# Population dynamics of Amazonian floodplain forest species support spatial variation on genetic diversity but not range expansions through time

**DOI:** 10.1101/2021.09.13.460077

**Authors:** Gregory Thom, Camila C. Ribas, Eduardo Shultz, Alexandre Aleixo, Cristina Y. Miyaki

## Abstract

**Aim:** We tested if historical demographic changes of populations occurring on the floodplains of a major Amazon Basin tributary could be associated with range expansions from upper and middle sections of the river, following the establishment of widespread river-created environments during the Late Pleistocene and Holocene.

**Location:** Solimoes River, Western Amazon, South America

**Taxon:** *Myrmoborus lugubris, Thamnophilus cryptoleucus* and *Myrmotherula assimilis* **Methods**: We analyzed thousands of UltraConserved Elements to explore spatial patterns of genetic diversity and connectivity between individuals. Range expansions were tested with alternative methods. We quantified habitat preference for the analyzed species in order to test if the occupation of dynamic habitats could predict spatial patterns of genetic diversity.

**Results:** Our study did not support shared population range expansions related to historical regionalized changes in habitat availability. We found considerable variation in the spatial distribution of the genetic diversity between studied taxa, and that species with higher levels of specialization to dynamic environments have a more heterogeneous distribution of genetic diversity and reduced levels of gene flow across space.

**Main conclusions:** Our results suggest that demographic expansions along the Solimões River might be linked to geographic homogeneous oscillation in the distribution of floodplain environments, promoting effective population size changes but not range expansion. We found that habitat specificity might be a good predictor of population connectivity along the Amazonian floodplains.

## Introduction

Understanding the factors that shape the spatial distribution of species is a long-standing challenge in comparative biology that is attributable to the complex interaction of environmental and species trait variables. Over evolutionary time scales the geographic distribution of populations might be affected by physiographic processes (Silva et al., 2019; Peter et al., 2020), climatic oscillations (Hewitt, 2000; Raposo do Amaral et al., 2018; Musher et al., 2020), biotic interactions (Araújo & Luoto, 2007; Wisz et al., 2013), and trait evolution (Alves et al., 2019), driving dispersion, extinction, and cladogenesis. Although processes such as the appearance of novel traits, interspecific competition, and colonization of new environments might lead to a single burst of range size change, in many cases species distributions are dynamic over time, oscillating between periods of range contraction followed by expansion (Davis & Shaw, 2001; Taberlet & Cheddadi, 2002; Carnaval & Moritz, 2008). Range expansions are expected to be accompanied by a rapid increase in the number of individuals, and strong spatial directionality (Excoffier et al., 2009). At the expansion front, successive founder events lead to a fast differentiation from the source, producing a characteristic demographic signature that is distinguishable from non-spatial population growth (Excoffier et al., 2009; François et al., 2010; Peter & Slatkin, 2013; Alvarado-Serrano & Hickerson, 2018). By tracking the spatial dynamics of populations one can locate zones of climatic stability (Potter et al., 2016), track sources of population invasions (Fischer et al., 2017), and understand how genetic diversity was affected by past environmental changes (Reid et al., 2019), allowing for more robust predictions to explore classical diversification hypotheses.

One of the main processes used by phylogeographic approaches to explain population size changes through time is the variation on available habitat distribution due to vegetation response to historical climate change, as described by the refugia hypothesis (Haffer, 1969, 1997; Hewitt, 2004). This mechanism suggests that populations tend to be isolated in “islands” of optimum habitats during glacial (e.g., lowland forest taxa) or interglacial (e.g., montane forest taxa) periods followed by population expansion and secondary contact as habitats expand. In the Neotropics, the region for which the refugia hypothesis was initially proposed (Haffer, 1969), the effects of past climatic oscillations have been used to explain patterns of genetic diversity in multiple environments, including the Atlantic Forest (Carnaval et al., 2009; Amaral et al., 2018; Silva et al., 2019), Amazonia (Thom et al., 2020b), Caatinga (Gehara et al., 2017), the Chaco dry forests (Camps et al., 2018) and the Andes/Patagonia (Cosacov et al., 2010). However, few studies have sought to understand if demographic patterns are linked to range size change (Camargo et al., 2013; Guarnizo et al., 2016; Baranzelli et al., 2017). By testing models including range expansions, one can explicitly explore alternative hypotheses regarding the location of putative stable habitats through time, providing a better understanding of the historical dynamic of specific environments (Potter et al., 2016; He et al., 2017; Reid et al., 2019).

In the Amazon floodplains, historical variation in precipitation and global sea-level affected sedimentation dynamics of the main rivers (Latrubesse & Franzinelli, 2005; Irion et al., 2009; Soares et al., 2010; Pupim et al., 2019), potentially shaping the distribution and connectivity of populations occurring on river-created environments (Aleixo, 2006; Choueri et al., 2017; Thom et al., 2020b). Phylogeographic studies have reported demographic expansions for populations of birds specialized in river islands, suggesting widespread habitat alterations across the entire Amazon river system (Aleixo, 2006; Thom et al., 2020b). Historical oscillations in the availability of floodplain environments tend to be more pronounced in regions where large tributaries with distinct sediment loads meet (Gualtieri et al., 2018). For instance, the confluence of the Negro (low sediment load) and Solimoes rivers (high sediment load) leads to an increase in the water discharge and dilution of sediment concentration of the main Amazon river channel, promoting erosion and sediment bypass (Thom et al., 2020b). The current dynamics between Negro and Solimoes rivers prevent the building of long-lasting substrates for flooded habitats downstream of the confluence for approximately 150 km, until the high sediment input of the Madeira River, which restores suitable conditions for the development of large and stable floodplains in the main channel of the Amazon river (Filizola & Guyot, 2009). Recent phylogeographic studies suggest that the central portion of the Amazon Basin, between the Negro and Madeira rivers, is indeed a suture zone, partially or completely isolating recently diverged populations, mostly restricted to Solimoes, Negro, Madeira, and lower Amazonas rivers (Choueri et al., 2017; Thom et al., 2018, 2020b). This spatial congruence indicates that fluvial sedimentation dynamics may directly influence the connectivity of species occurring in river-created environments along Amazonian floodplains through time.

The historical interaction between the sediment-poor Negro River and the sediment-rich Solimões-Amazonas main channel, mediated by past oscillations in precipitation and sea level, may have been dynamic over time, shaping the availability of flooded habitats in central Amazonia. For example, the Negro River is currently impacted by a damming effect caused by the larger water input from the Solimões-Amazonas River, maintaining a ria lake at its lower section, which prevents island formation and creates a barrier for the biota associated with river-created habitats (Choueri et al., 2017). Similarly, past increases in the Negro River discharge could have periodically affected the flooding pulse of the Solimões (Irion et al., 2009; Passos et al., 2020), causing suppression of seasonally flooded habitats from its lower and middle sections (Latrubesse & Franzinelli, 2005; Soares et al., 2010; Passos et al., 2020). Under this scenario, we expect seasonally flooded habitats in the upper Solimões River to be historically more stable than areas closer to the Negro River confluence (lower Solimoes River). Hence, the demographic expansions detected by previous studies for populations of birds (Thom et al., 2020b) would be linked to recent range expansions from the upper/middle towards the lower sections of the Solimoes River. This demographic process should lead to higher genetic diversity on upriver populations, and if this was a pervasive mechanism, we expect to detect the same pattern in co-occurring populations that occupy similar habitats. Alternatively, population demographic expansions could be related to geographic homogeneous oscillation in habitat availability (e.g., oscillations in the carrying capacity of the environments). Under this scenario, we expect the genetic diversity to be mostly explained by the geographic distance between individuals (Isolation by distance), with a relatively homogeneous distribution of the genetic diversity across the whole extension of the Solimoes River, or governed by species-specific characteristics. In the latter, species would have variable patterns across space, potentially associated with habitat specificity.

The Amazon floodplains are an intricate mosaic of microhabitats that roughly segregate with the sedimentary dynamic of the rivers and the geographic distance from the main channel (Junk et al., 2012, 2015). Areas next to the margins or on islands of rivers with high sedimentary budgets as the Solimoes, tend to be more dynamic over time, keeping vegetation in early successional stages of regeneration due to the constant erosion and sedimentation. Given the high diversity of microhabitats along the floodplains, we might expect that species with higher dependence on more dynamic environments should have been more affected by historical changes on the floodplains, having a more heterogeneous spatial distribution of genetic diversity. In this case, higher association to more dynamic habitats should predict spatial levels of genetic diversity.

We explored the levels of connectivity and the spatial distribution of the genetic diversity of three antbird species, *Myrmoborus lugubris, Thamnophilus cryptoleucus*, and *Myrmotherula assimilis* with populations restricted to the Solimões River, testing if the population demographic expansions reported in a previous study (Thom et al., 2020b) are related to range expansions following a period of isolation in the upper/middle course of this river. We reanalyzed the data published by Thom and collaborators (2020b) using a set of methods designed to track population range expansions and explore how genetic diversity is distributed across space. Our results refute the idea of congruent range shifts related to historical regionalized changes in habitat availability. Instead, our data suggest that demographic expansions along the Solimões floodplains might be associated with variation in habitat availability across the entire range of the species, promoting effective population size changes but not range expansion. We found that species with higher levels of specialization to dynamic environments have a more heterogeneous distribution of genetic diversity and reduced levels of gene flow across space, indicating that habitat specificity might be a good predictor of population connectivity along the Amazonian floodplains.

## Material and Methods

We selected samples from the Solimões River populations of *Myrmoborus lugubris* (N = 31), *Thamnophilus cryptoleucus* (N = 20), and *Myrmotherula assimilis* (N = 16) that were previously analyzed by Thom et. al (2020b; Figure 1; Table S1). Genome-wide variation was obtained through the sequence capture of Ultraconserved Elements (UCEs) using a probe set targeting 2,312 UCE loci and 97 additional probes targeting exons commonly used in Avian phylogenomic studies (Hackett et al., 2008; Kimball et al., 2009; Faircloth et al., 2012). Raw sequences were downloaded from the Sequence Read Archive (SRA; BioProject ID: PRJNA595086; SubmissionID: SUB6206537; www.ncbi.nlm.nih.gov/bioproject/595086). We used Trinity 2.4 (Grabherr et al., 2011) to assemble *de novo* reads into contigs. The contigs were matched to the probes with LASTZ (http://www.bx.psu.edu/~rsharris/lastz/) using “match_contigs_to_probes.py” from PHYLUCE (Faircloth et al., 2012; Faircloth, 2016). Contigs that did not align to probe sequences and those that matched multiple loci were removed. The resulting fasta files from the different individuals were aligned in MAFFT v7.475 (Katoh & Standley, 2013), allowing for missing individuals and without trimming long ragged-ends. These long ragged-ends were then trimmed by applying a threshold of 50% of missing sequences among individuals with TrimAl v1.4.rev15 (Capella-Gutierrez et al., 2009). The longest sequence without indels of each locus was selected as a reference for the following steps. First, reads of each individual were aligned to the reference allowing 4 mismatches per read using BWA v0.7.17-r1188 (Li & Durbin, 2009). The sam files obtained were converted to bam format with Samtools v1.10 (Li et al., 2009). Reads were trimmed to match the reference using CleanSam.jar, reassigned to groups with AddOrReplaceReadGroups.jar, and duplicated reads were identified with Markduplicates.jar from PICARD v.2.0.1 (Broad Institute, Cambridge, MA; http://broadinstitute.github.io/picard/). We merged individual bam files into a single bam file of all samples with MergeSamFiles.jar from PICARD. With GATK v3.6 (McKenna et al., 2010) we realigned all reads, identified indels (RealignerTargetCreator; IndelRealigner), and called SNPs hard-masking low-quality bases (< Q30) with UnifiedGenotyper and VariantAnnotator. We obtained raw vcf files for each species that were filtered for a minimum read depth of > 8 using VCFTOOLS v0.1.15 (Danecek et al., 2011). Loci with any site with heterozygosity higher than 0.75 were excluded. Finally, we randomly selected one SNP per locus, excluding sites with missing data, resulting in a complete SNP matrix.

**Figure 1:**
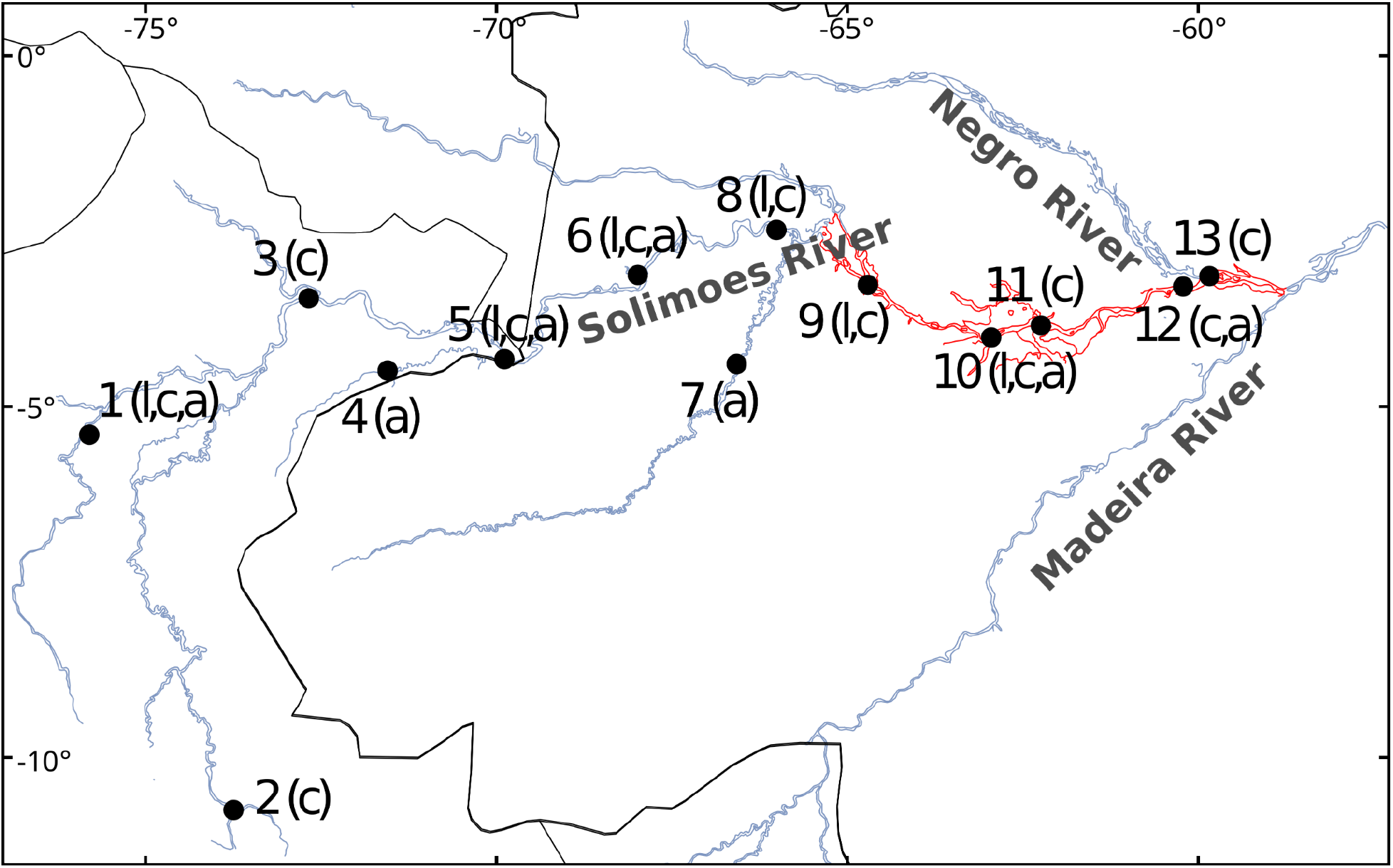
Geographic distribution of sampled individuals of *Myrmoborus lugubris* (l), *Thamnophilus cryptoleucus* (c), and *Myrmotherula assimilis* (a) across the Solimões River Basin. Numbers in the map represent localities as in Table S1. In red is the putative zone of unstable habitats following the range expansion hypothesis.

### Genetic Structure

We tested the best-fit number of ancestral populations (k) for each species complex and clustered individuals to populations in sNMF v1.2 (Frichot et al., 2014) by applying a sparse non-negative matrix factorization to compute least-square estimates of ancestry coefficients. We explored values of k between 1 and 8 and performed 100 replicates for each value with an alpha regularization parameter of 100. Given that populations occurring in linear and continuous environments such as rivers are more subjected to the effects of the geographic distance among individuals, and that IBD can generate discrete genetic clusters (Thomaz et al., 2016), we tested genetic structure controlling for the effects of IBD in the R package conStruct v1.0.3 (Bradburd et al., 2018). ConStruct simultaneously estimate continuous and discrete patterns of population structure by assuming a rate of decay in the relatedness among individuals as a function of the geographic distance. Thus, discrete groups are only assumed when genetic variation significantly deviates from an IBD scenario. We ran models assuming the same number of layers as the number of ancestral populations obtained in the best sNMF model and calculated the relative layer contribution with the function “calculate.layer.contribution”, in order to observe if the genetic structure inferred by sNMF could be explained exclusively by IBD. For each conStruct run, we performed 50,000 iterations discarding the first 50% as burn-in. We calculated the geographic distance matrices for each species following river connectivity.

### Isolation by distance, effective diversity, and gene flow

We identified spatial variation in gene flow among localities and tested deviations from isolation by distance (IBD) scenario for each species with EEMS (Petkova et al., 2016). EEMS estimates the effective migration surface among drawn demes mapping genetic differentiation based on a spatially explicit approach by using a Markov Chain Monte Carlo (MCMC) to estimate effective demographic parameters given the observed genetic dissimilarity between individuals. Euclidian genetic dissimilarity matrices between individuals for each species complex were generated in ADEGENET v2.1.3 (Jombart & Ahmed, 2011). Habitat polygons were produced based on the geographic distribution of each species complex and 300 demes were distributed over the habitat area. Each MCMC run was performed for 30×10^6^ generations with the first 5×10^6^ generations excluded as burn-in. Maps were generated using additional features of the EEMS R package. We calculated the inbreeding coefficient (F) per individual in VCFTOOLS and interpolated the obtained values across the species distribution in QGIS (https://www.qgis.org/). Additionally, to test IBD we performed a Mantel correlation test between genetic and geographic distances in ADEGENET with 999 replicates.

### Geographic range expansion

We tested for population range expansion using two alternative approaches, first estimating the directionality index using rangeExpansion v0.0.0.9 in R (Peter & Slatkin, 2013, 2015) and second, estimating the geographic spectrum of shared alleles in the GSSA v0.0 program (Alvarado-Serrano & Hickerson, 2018). The rangeExpansion applies a founder effect algorithm assuming a stepping stone model of population expansion from a single location, testing the strength of the spatial expansion and the most likely origin of the founder effect, measuring the effective founder distance. The effective founder distance represents the size of the deme in which the effective population size is reduced by 1% during the founder event occurring in the expansion front. Thus, low values for the effective founder distance suggest a strong founder effect. For this approach, in order to calculate the derived state of each SNP, we assumed as outgroup one individual from a closely related population (Thom et al., 2020b).

The GSSA (Alvarado-Serrano & Hickerson, 2018) captures the range expansion signal by summarizing the shared co-ancestry between sampled localities. This statistic consists of estimating the absolute number of minor alleles shared between sampled localities, taking into account the geographic distance between localities. Given that the amount of genetic drift increases with the distance to the founder event (source of the expansion), the obtained per locality histograms contain information regarding the position of a given locality with respect to the source of the geographic expansion. Some advantages of the GSSA are that no outgroup needs to be used and it can handle less geographic dense sampling when compared to the rangeExpansion method. GSSA implementation allows for the user to provide a geographic distance matrix, instead of assuming linear distances from geographic coordinates. Given the intimate relationship of the studied species with the floodplains, we calculated the geographic distance matrixes for each species following the river’s connectivity. The information contained in the shape of the GSSA histogram was summarized using Harpending’s raggedness index (Harpending, 1994) and the obtained values per locality were interpolated across the distribution of each species in QGIS. We expect that locations further away from the expansion source share more genetic variants with nearby locations that were colonized through the same expansion route, and thus have higher values for the Harpending’s raggedness index. For both analyses, we selected a single individual per locality with the lowest amount of missing data.

### Habitat occupancy and river dynamism

To quantify differences in habitat occupation between the three species that could predict the spatial patterns of genetic diversity we used a metric of river dynamism extracted from the Occurrence Change Intensity Map available at https://global-surface-water.appspot.com/ (Pekel et al., 2016). This layer contains information on whether the water surface increased (e.g. erosion) or decreased (e.g. deposition of sediments) around bodies of water, and its intensity based on both the intra and inter-annual variability and changes, extracted from Landsat satellite images between 1984 and 2019. On this layer, values range from −100 to 100, where negative values represent a loss of water surface, positive values represent an increase of water surface, and zero represent stable water surfaces. To calculate the dynamicity of a radius around occurrence records for the studied species, we: 1) converted negative values to positive, so both gain and loss represent how much a given pixel has changed over time; 2) drew a buffer of 2km around every stable body of water, removing areas that were permanently water across the analyzed time section; 3) used random drawn points and occurrence records for the three species to calculate average values of dynamicity using a 1000 meters radius around the coordinates. To create the random distribution we draw 1000 points across the floodplains of the Solimoes Basin. We plotted the values for dynamicity and calculated pairwise Wilcoxon tests in R, to explore if values extracted from random points and each species statistically differ. Spatial layers were processed using the raster v3.4 in R (Hijmans & van Etten, 2013). Occurrence records were obtained from video, photos, and sound recordings from Macaulay Library (www.macaulaylibrary.org) and Xeno-canto (www.xeno-canto.org), and deposited specimens on scientific collections (Table S2). Each record was individually checked, and records with a generic locality description or low-resolution coordinates were excluded. Occurrences at the same locality and date were counted only once. Geographic coordinates outside the 2km buffer were moved to the closest pixel within the buffer.

## Results

Our bioinformatics pipeline produced raw vcf files with 5,602 (16.8 mean depth; 2.8 sites per loci), 10,624 (28.4 mean depth; 4.7 sites per loci), and 10,283 (19.9 mean depth; 7.1 sites per loci) variant sites for *M. lugubris, T. cryptoleucus*, and *M. assimilis*, respectively. After randomly selecting one SNP per UCE loci we obtained matrices with 1,476, 1,805, and 1,776 SNPs for *M. lugubris, T. cryptoleucus*, and *M. assimilis*, respectively. The number of genetic clusters obtained with sNMF was partially concordant between species. For *M. lugubris* and *T. cryptoleucus*, the best value of masked cross-entropy was achieved with K=2 while for *M. assimilis* the best value suggested K=1 (Figure 2, S1). However, the results for K=2 for *M. assimilis* recovered a similar geographic structure as the ones observed in the other two species, with a gradual transition over the central portion of the Solimões basin (Figure 2). For *T. cryptoleucus* and *M. assimilis*, this transition in the ancestrality coefficients occurred between Tefé and Codajas (localities 9 to 11 in Figure 1) while for *M. lugubris* it occurred more to the west, at Santo Antonio do Ica (locality 6 in Figure 1). When accounting for geographic distance with conStruct, assuming 2 layers, the relative contribution of the second layer was less than 1% and without geographic correspondence in all three species. These results suggest that the genetic structure obtained with sNMF is better explained by isolation by distance (IBD).

**Figure 2:**
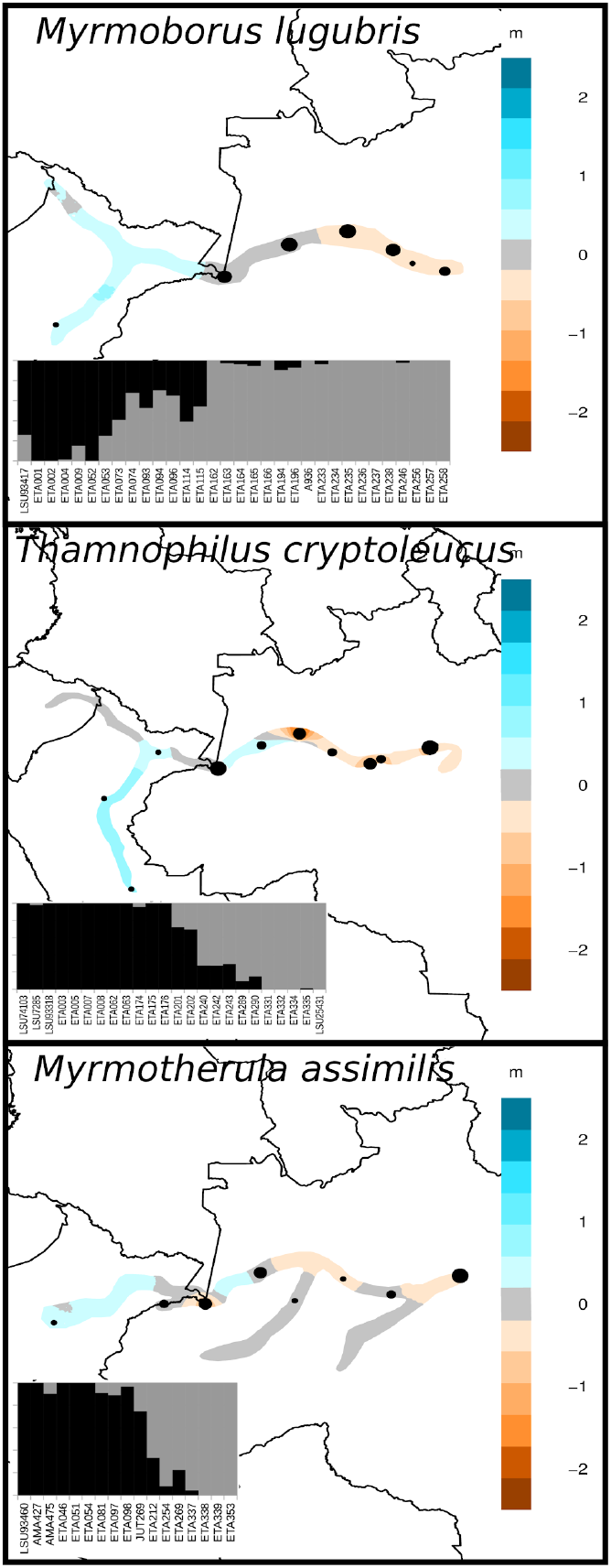
Estimated effective migration surface (EEMS) and population structure and individual coefficient of ancestry (bars) inferred with sNMF for the three species studied. Colors in the map represent a log10 scale from the average effective migration (gray). Codes below the bars in the admixture plot represent identification numbers of individuals as in Table S1. Note that for *M. assimilis*, K=2 was the model with the second-highest cross-entropy value.

The EEMS detected slightly lower effective migration than the average in the eastern Solimões basin, suggesting progressively more dissimilar individuals towards the east in all three species (Figure 2). However, the estimated effective diversity produced contrasting results between species. While for *M. lugubris* we detected lower diversity in eastern localities, for *T. cryptoleucus* there was considerably lower diversity in western localities, and for *M. assimilis* we found a relatively homogeneous distribution of genetic diversity (Figures S2, S3). The results for a Mantel test were congruent with EEMS analyses, significantly supporting an IBD scenario in all three species (p-value < 0.01; Mantel r statistics of 0.59, 0.45, and 0.74 for *M. lugubris, T. cryptoleucus*, and *M. assimilis*, respectively). The inbreeding coefficient indicated a similar scenario as reported by EEMS effective diversity results for *M. lugubris* and *M. assimilis*, and a homogeneous distribution for *T. cryptoleucus* (Figure S4).

The geographic expansion model was strongly supported over an equilibrium/IBD model in the rangeExpansion analysis only for *T. cryptoleucus* (p-value < 0.001; q = 0.0004; Table 1; Figure S5) indicating a geographic expansion from western Solimões. For *M. lugubris* and *M. assimilis* the equilibrium/IBD model could not be rejected (p-value of 36.17 and 0.86 for *M. lugubris* and *M. assimilis*, respectively). The GSSA estimations support a geographic directionality for the raggedness index in *M. lugubris*, with lower values in the eastern and higher values in the western Solimões as suggested by the rangeExpansion (despite the non-significant p-value; Figure 3). For *T. cryptoleucus*, despite the significant results in the rangeExpansion, there is no geographic signal supporting a range expansion using the GSSA.

**Figure 3:**
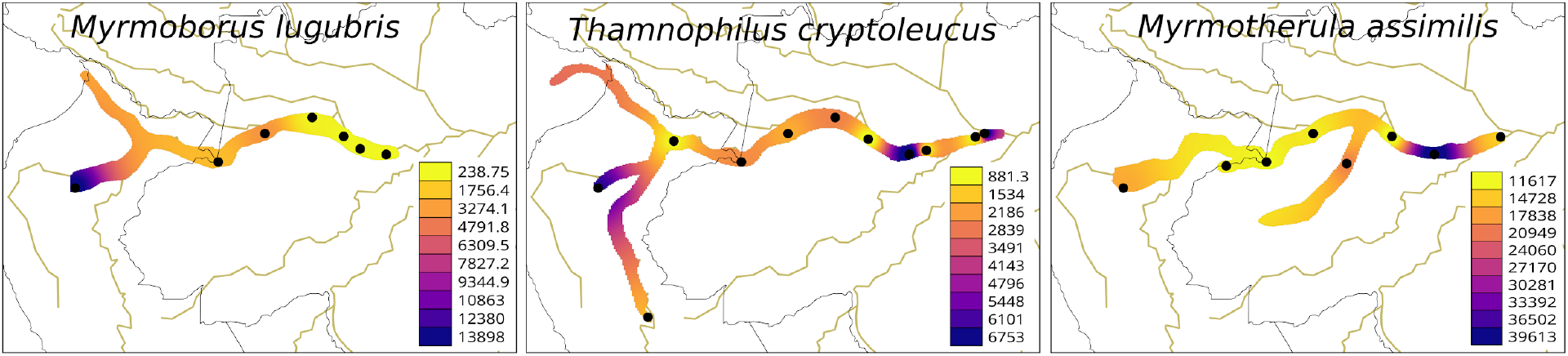
Interpolated Harpending’s raggedness index estimated from the geographic spectrum of shared alleles (GSSA) histograms. Lower values represent the most likely source of a serial range expansion.

**Table 1:** Summary of results obtained with rangeExpansion for three species of birds occurring across the Solimoes River.

| Parameters | *M. lugubris* | *T. cryptoleucus* | *M. assimilis* |
| --- | --- | --- | --- |
| longitude | -62.95 | -73.22144 | -75.64357 |
| latitude | -4.021451 | -9.999475 | -3148 |
| r1 | 0.999 | 0.999 | 0.999 |
| r10 | 0.996 | 0.991 | 0.997 |
| r100 | 0.964 | 0.918 | 0.975 |
| d1 | 27.19 | 11.45 | 40.80 |
| r<sup>2</sup> | 0.45 | 0.89 | 0.31 |
| p-value | 36.17 | 2.10e-23 | 11.26 |

r1, r10, and r100 - decrease in diversity at 1, 10, and 100 kilometers of distance, respectively; d1 – effective founder distance, which represents the deme size in km for the effective population size to be reduced by 1% in the expansion front; r<sup>2</sup> and p-value - correlation coefficient and correlation p-value for the most likely origin, respectively.

Data processing of occurrence records produced 60, 92, and 44 points for *M. lugubris, T. cryptoleucus*, and *M. assimilis*, respectively. The pairwise Wilcoxon tests indicated that all three species occupy areas that are more dynamic than randomly drawn points, with *M. lugubris* and *T. cryptoleucus* occurring in areas significantly more dynamic than *M. assimilis* (Figure 4).

**Figure 4:**
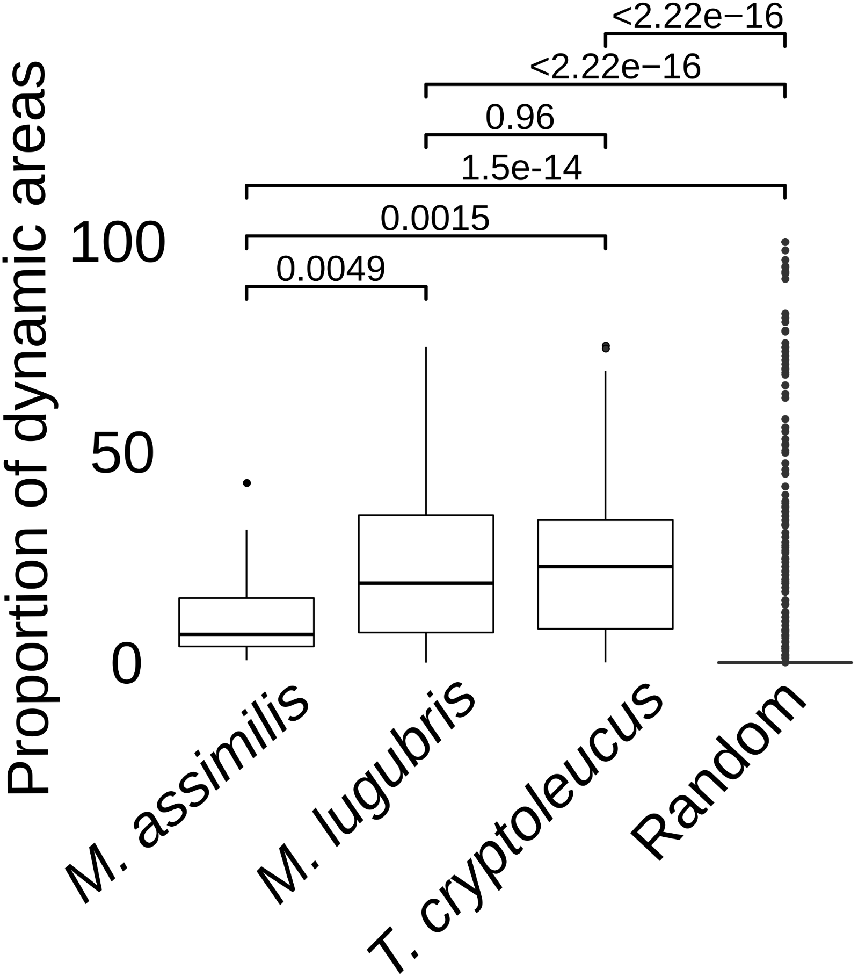
Average river dynamism on 1000 meters radius around randomly drawn points and occurrence records for *M. assimilis, M. lugubris*, and *T. cryptoleucus*. River dynamism was calculated based on the “Water Occurrence Change’’ layer available at https://global-surface-water.appspot.com/. Higher values represent higher river dynamicity. The pairwise comparisons show p-values for Wilcoxon tests between species and random points.

## Discussion

### Demographic histories are not linked to a shared pattern of range expansion

We used the genetic variation within thousands of Ultraconserved Elements to explore spatial patterns of genetic diversity in three species of antbirds that inhabit river-created environments along the Solimões River. Previous studies suggested that populations of these three species experienced synchronic demographic expansions in the last 0.2 million years ago (Thom et al., 2020b). Here we tested if demographic expansions were linked to geographic range expansions associated with historical alteration in the distribution of river-created environments (Latrubesse & Franzinelli, 2005; Irion et al., 2009; Soares et al., 2010; Govin et al., 2013; Pupim et al., 2019). Our results supported that only one species, *T. cryptoleucus*, had a significant range expansion with geographic origin in the western Solimoes River. The spatial distribution of the genetic diversity reported here refutes historical events triggering common regionalized changes in population ranges along the Solimoes River. Our results suggest that despite the high dynamism of floodplain environments over time, significant regional suppression of seasonally flooded habitats was probably restricted to specific portions of the floodplains. For example, the central portion of the Amazon, where the sedimentary basin of the Solimões River flows into the eastern Amazon basin, delimits the distribution of many floodplain lineages and is a suture zone splitting populations and species with distinct levels of genetic structure and gene flow (Farias & Hrbek, 2008; Albernaz et al., 2012; Zimmer & Isler, 2016; Thom et al., 2020b). Hence, the demographic expansions reported by Thom and collaborators (2020) might be related to homogeneous oscillations in habitat availability across the entire geographic distribution of the populations, promoting population size changes but not directional range contractions and expansions.

### Genetic diversity is heterogeneously distributed across the Solimoes floodplains

We observed species-specific patterns for the spatial distribution of genetic diversity along the Solimões River, potentially associated with variation in the occupancy of dynamic habitats (Figures 2, S2, S3, S4). The antbird species studied here are intimately related to the river edge forests that are mostly distributed over islands of large rivers (Remsen & Parker, 1983; Junk et al., 2011). Given the low dispersal capacity of the studied species (Zimmer & Isler, 2016) and the historical dynamism of river islands, one might expect a shared pattern of genetic diversity between species. However, our data do not support this scenario, with the three species having different patterns of the spatial distribution of their genetic diversity. Despite occupying similar environments, our data suggest that species differ in their levels of association to dynamic habitats. We found that species occurring in significantly more dynamic areas have higher inbreeding coefficients, more heterogeneous spatial distribution of genetic diversity, and more variable levels of effective gene flow across space. These results indicated that habitat specificity might predict the spatial connectivity of floodplain species. Despite the congruency in historical demography, reported in previous studies (Thom et al., 2020b), our results suggest a pronounced effect of intrinsic ecological traits leading to distinct dispersal capacities affecting the spatial distribution of the genetic diversity within populations.

Heterogeneous patterns of genetic diversity have been reported along the Amazonian floodplains (Choueri et al., 2017; Jardim de Queiroz et al., 2017; dos Anjos Oliveira et al., 2019; Farias et al., 2019). Choueri and collaborators (2017) studying landscape genetics of river island specialist birds found that genetic diversity over Negro River archipelagos is not homogeneously distributed and that the temporal dynamics of formation and disappearance of islands shaped the genetic structure and gene flow between populations. Additionally, these authors suggest that the increase in sediment accumulation and island formation during the Holocene expanded connectivity between archipelagos (Latrubesse & Franzinelli, 2005; Latrubesse & Stevaux, 2015; Choueri et al., 2017). However, in contrast to the demography of Solimões River populations, only slight signals of expansion were detected for the Negro River populations, corroborating the idea that rivers with black and clear water (sediment-poor) are less dynamic over time than white-water rivers (sediment-rich rivers), such as the Solimões (Thom et al., 2020b).

The patterns of genetic structure we detected with sNMF (K=2) were likely a product of the strong effects of IBD in linear distributions, as observed in the conStruct analyses, which suggested a minor contribution of a second layer (K) to explain the genetic structure. These results are in agreement with simulation-based, and empirical studies reporting the high effect of IBD in the linear distribution of river systems, including the formation of differentiated lineages (Selkoe et al., 2015; Thomaz et al., 2016; Farias et al., 2019). However, studies have suggested that the riverway distance plays a minor role in structuring genetic diversity of floodplain populations across the entire Amazonian Basin, suggesting the more pronounced effect of other environmental features (Hrbek et al., 2005; dos Anjos Oliveira et al., 2019; Thom et al., 2020b). Our results reinforce the idea that IBD has to be taken into account when testing for genetic structure in continuous populations on the risk of assuming spurious discrete populations in downstream population genetics analyses.

### Future directions to explore spatial patterns of genetic diversity across the Amazonian floodplains

Our study provided new insights on the spatial dynamics of floodplain forest environments over time, refuting a pattern of common range expansion for co-occurring species. However, our data is limited in the sense that more sampled localities could improve our power in detecting range size changes. Although studies have reported robust results with the rangeExpansion method (Peter & Slatkin, 2013) sampling a single diploid individual for 10 localities and 1,500 SNPs, which is a data set similar to ours, the ideal data set lies around 20 diploid individuals (1 per locality) and 7,000 SNPs (Peter & Slatkin, 2013, 2015; Potter et al., 2016). An important factor that might hinder our ability to detect if a range expansion occurred is the time since populations reached a stationary condition (Alvarado-Serrano & Hickerson, 2018). All three species we analyzed have extensive distributions across the Solimões River, and might already have reached isolation by distance equilibrium, eroding the signal for directional range expansions. Recent evidence has supported that secondary contact and gene flow are pervasive processes across the floodplains, although controlled by varying degrees of connectivity through time (Thom et al., 2018, 2020b). The methods we applied do not incorporate the effects of gene flow from unsampled populations, which could inflate genetic diversity in eastern Solimões localities, masking the genomic signal of range expansions. The inferred range expansion in *T. cryptoleucus* pointed to an interesting scenario, since this taxon is currently in contact with *T. nigrocinereus tschudi* from the Madeira River, suggesting diversification in allopatry followed by range expansion and recent secondary contact. These results suggest that the approach we used might be robust to some level of gene flow from distinct populations. Future studies aiming to explore the spatial demographic dynamics of floodplain populations should target widespread geographic distribution of samples (e.g. one individual per locality and many localities), and more genetic markers, obtained with reduced representation techniques such as RADseq (Andrews et al., 2016), or whole-genome sequencing (Ellegren, 2014). Additionally, spatial explicit simulations of demographic histories coupled with supervised machine learning classification algorithms could improve our ability to test range size changes through time with a reduced number of samples (Battey et al. 2020; Reid et al. 2019).

Ecological niche modeling has been used to track the current and paleo distribution of various Amazonian organisms (Prates et al., 2016; dos Anjos Oliveira et al., 2019; Silva et al., 2019). However, the intimate relationship of floodplain species with microhabitats linked to the sedimentation dynamics of the rivers, probably cannot be captured by interpolated bioclimatic variables (e.g. Worldclim). Meaningful variables that could describe the sedimentation process and detailed distribution of floodplain habitats with a good resolution for the entire Basin are becoming available (Pekel et al., 2016; Fassoni-Andrade & Paiva, 2019) and might provide new insights on how the current sediment budget of rivers and vegetation dynamics shape patterns of genetic diversity. Amazonia is currently at the center of the debate about the impact of hydroelectric dams on biodiversity and the sedimentary budget of rivers (Latrubesse et al., 2017). Understanding how the genetic diversity (dos Anjos Oliveira et al., 2019) and species richness (Laranjeiras et al., 2021) might be associated with river dynamism and productivity is paramount to predict future impacts caused by anthropic activities.

## Acknowledgments

We thank the curatorial staff of the Museu Paraense Emílio Goeldi (MPEG), Instituto Nacional de Pesquisas da Amazônia (INPA), Laboratório de Genética e Evolução Molecular de Aves (LGEMA) for loaning tissue samples under their care. We thank members of the Smith lab, Hickerson lab, and Carnaval lab: Brian Smith, Kaiya L. Provost, Lucas R. Moreira, Vivien Chua, Jon Merwin, Ana C. Carnaval, Connor French, Andrea Paz, Kathryn Mercier, Rilquer Mascarenhas, Lidia Martins, Michael Hickerson, Isaac Overcast, Alexander Xue. We thank L. E. Araujo-Silva for the partnership during field expeditions. We thank Oscar Johnson and Rob Brumfield for sharing UCE sequences. We thank the Research Center on Biodiversity and Computing (BioComp) of the Universidade de São Paulo (USP), supported by the USP Provost’s Office for Research. This study was co-funded by FAPESP (BIOTA, 2012/50260-6 and 2013/50297-0), NSF (DOB 1343578), NASA, CNPq (310593/2009-3, 574008/2008-0, 563236/2010-8, and 471342/2011-4), PEER/USAID (AID-OAA-A-11-00012), and FAPESPA (ICAAF 023/2011). G.T. was granted by CAPES and then FAPESP scholarships (2014/00113-2, 2015/12551-7, 2018/17869-3, and 2017/25720-7). A.A., C.C.R., and C.M. are supported by CNPq research productivity fellowships (310880/2012-2, 308927/2016-8, and 3062204/2019-3).

## Biosketch

**Gregory Thom** is broadly interested in comparative genomics, testing the major factors impacting population differentiation over historical timescales. His overriding goal is to unearth the mechanisms driving Neotropical bird evolution by using genomics and advanced computational biology.

## Author contributions

G.T. conceived the study and performed bioinformatics and phylogeographic analyses. G.T., C.C.R., and C.M. wrote the paper with input from A.A. and ES. GT and ES performed the river dynamicity analyses. All authors reviewed and approved the final version of the manuscript before submission.

